# Accuracy of a machine learning method based on structural and locational information from AlphaFold2 for predicting the pathogenicity of *TARDBP* and *FUS* gene variants in ALS

**DOI:** 10.1101/2022.07.07.499092

**Authors:** Yuya Hatano, Tomohiko Ishihara, Osamu Onodera

## Abstract

**Background:** In the sporadic form of amyotrophic lateral sclerosis (ALS), the pathogenicity of rare variants in the causative genes characterizing the familial form remains largely unknown. To predict the pathogenicity of such variants, *in silico* analysis is commonly used. In some cases of ALS, the gene mutations are concentrated in specific regions, and the resulting alterations in protein structure are thought to significantly affect pathogenicity. However, existing methods have not taken this issue into account. To address this, we have developed a technique termed MOVA (method for evaluating the pathogenicity of missense variants using AlphaFold2), which applies positional information for structural variants predicted by AlphaFold2. Here we examined the utility of MOVA for analysis of several causative genes of ALS.

**Methods:** We analyzed variants of six ALS-related genes (*TARDBP*, *FUS*, *SETX*, *TBK1*, *OPTN*, and *SOD1*) and classified them as pathogenic or neutral. For each gene, the features of the variants, including their positions in the 3D structure predicted by AlphaFold2, were entered into a random forest algorithm and evaluated by leave-one-out cross-validation. We compared how accurately MOVA was able to classify the pathogenic and neutral mutation variants.

**Results:** MOVA yielded useful results (AUC ≥0.70 for 3 (*TARDBP* 0.755, *FUS* 0.844, and *SOD1* 0.787) of the 6 genes) and was particularly useful for genes where pathogenic mutations were concentrated at specific sites (*TARDBP*, *FUS*).

**Conclusions:** MOVA is useful for predicting the virulence of rare variants of ALS-causing genes in which mutations are concentrated at specific structural sites.

## Introduction

Rare variants in the causative genes of familial amyotrophic lateral sclerosis (FALS) are found in 10-30% of cases of sporadic ALS (SALS).[1, 2] However, the pathological significance of such variants occurring only in SALS is largely unknown. The pathogenicity of these rare variants must be validated using cultured cells, animal models, and samples obtained from patients. However, this type of validation is generally difficult to perform. Instead, *in silico* analytical methods can be used to predict pathogenicity.[3–7]

*In silico* analysis focuses primarily on evolutionary conservation of gene and amino acid similarity; the most commonly used algorithm is PolyPhen-2, which is a machine learning approach employing eight variables based on nucleotide sequences and variables predicted from known three-dimensional (3D) structures.[3] REVEL and CADD are methods that integrate several *in silico* analysis methods, including PolyPhen-2.[4, 5] EVE, a machine-learned method utilizing evolutionary conservation of sequences, has also been reported to predict the pathogenicity of rare variants without supervised data.[6] Neither method alone is recommended for identifying pathogenicity. Guidelines for genetic diagnosis from the American College of Medical Genetics and Genomics (ACMG) and the Association for Molecular Pathology (AMP) recommend that these analytical methods should be used only when multiple methods predict that the variant is deleterious.[8] Therefore, development of a new *in silico* approach from a new perspective would be desirable.

One factor not often considered in existing *in silico* analysis methods is the location of the variant and the 3D structure of the associated region.[7] In the ALS-associated genes *TARDBP* and *FUS*, mutations are concentrated in a particular region.[9, 10] These regions are also structurally characteristic and are assumed to undergo facilitated aggregation. If positional information and pathogenicity related to the structure of these regions can be taken into account, more accurate estimation of pathogenicity might be possible. Indeed, the ACMG guidelines recommend that even if a variant is located in a mutational hotspot or in an important functional domain, it is a factor that would support the pathogenicity of the variant. Therefore, the accuracy of pathogenicity prediction would be improved by taking into account positional information about the variants. However, it has been difficult to utilize this factor because of the limited number of proteins whose 3D structures have been clarified.[7]

Recently, AlphaFold2 was developed as a method for prediction of protein 3D structures *in silico* with high accuracy.[11] This method can predict the structure of a protein even in the absence of a similar protein whose structure is already known.[12] Therefore, addition of structural information using AlphaFold2 would be expected to improve the accuracy of pathogenicity prediction. In fact, AlphScore has been shown to predict the pathogenicity of rare variants using this factor,[7] and when combined with REVEL, CADD, and DEOGEN2, it improved the accuracy of each program.[7] These results suggest that structural information from AlphaFold2 would be useful for pathogenicity prediction.

Here we developed an approach termed MOVA (a method for evaluating the pathogenicity of missense variants using AlphaFold2), which applies positional information for variants based on the 3D structure predicted by AlphaFold2, and machine-learns the pathogenicity of variants for each gene. We investigated the usefulness of MOVA for predicting the pathogenicity of rare ALS-causative gene variants.

## Materials and Methods

### Gene set

Among the known ALS-causative genes, six genes (*TARDBP*, *FUS*, *SETX*, *TBK1*, *OPTN*, and *SOD1*) with a large number of pathogenic missense variants, were included in the analysis. The pathogenic variants were defined as indicated below.

### Data set

Variants of each gene listed in gnomAD v.3.1.2 or HGMD Professional 2022.1 were included in the analysis. Pathogenic variants (positive variants) were defined as variant class ‘DM’ with reported phenotypes including amyotrophic lateral sclerosis, frontotemporal dementia, or motor neuron disease in HGMD Professional 2022.1. Neutral variants (negative variants) were defined as variants recognized in gnomAD v.3.1.2 excluding variants defined as a class ‘DM’ or ‘DM?’ in HGMD Professional 2022.1.

### CADD, PolyPhen-2, EVE, REVEL and AlphScore

We evaluated the accuracy with which CADD, PolyPhen-2, EVE, REVEL, and AlphScore were able to discriminate between positive and negative variants in the existing datasets. PolyPhen-2 used HumDiv as the classifier model. Sensitivity (true-positive rate) versus 1-specificity (false-positive rate) at each threshold was plotted as a ROC curve, and the AUC was calculated. The specificity and sensitivity at each threshold were calculated using the prediction function of the ROCR package in R version 4.1.0, and the ROC curve plot and AUC were calculated using the performance function.

### MOVA: a method for determining variant pathogenicity using AlphaFold2

To construct MOVA, we used supervised machine learning with labeled examples attributed to positive or negative variants, and trained and examined each gene individually. The average x, y and z coordinates for each atom and the pLDDT (predicted local distance difference test) score at the site of the mutant amino acid residue of the protein in the pdb file of the Alphafold2 database,[13] and “the likelihood of an event in which the reference amino acid at the site of amino acid change remains the same – the likelihood of an event in which the reference amino acid is replaced by the alternative amino acid, as evaluated by BLOSUM62” were used as features. The training was performed using the random forest algorithm (Figure 1A).[14] The pLDDT is an estimate of how well the predictions match the experimental structure based on the local distance difference test Cα (lDDT-Cα), which is generated on a scale of 0-100 for each amino acid residue.[11] The model was constructed using the randomForest function of the randomForest package for the statistical software R version 4.1.0. The parameters were left at the default settings of the randomForest function.

**Figure 1.**
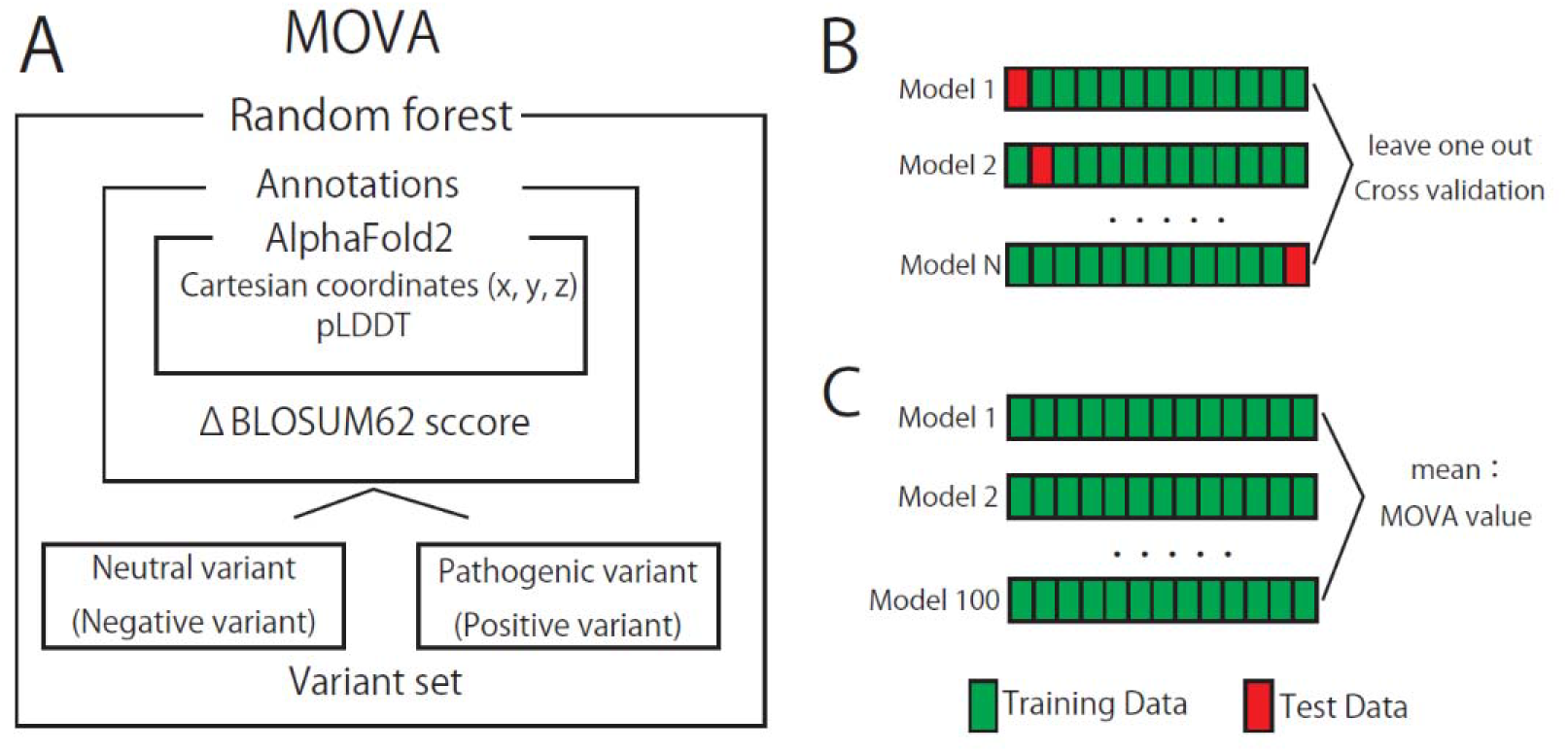
Work flowchart for MOVA. The x, y, z coordinates, and the plddt score for the amino acid residues at the substitution sites in the protein in the pdb file of the Alphafold2 database, and the ΔBLOSUM62 of the substituted amino acid residue, were used as parameters for random forest training (A). Only one variant was extracted from the sample group and used as test data, and the rest were used as training data to build the model. The predictions were calculated and validated on the test data. This was repeated for as many of the data as possible so that all variants were test data (B). The model was generated 100 times with all variants in the dataset as training data. The probability of each possible variant of the gene being pathogenic was predicted, and the average of the predictions was used as the MOVA value (C).

### MOVA model evaluation

In this analysis, the model was evaluated by leave-one-out cross-validation (Figure 1B). One fold was defined as construction of the model by extracting only one case from the sample group as test data and using the rest as training data. Predictions (looMV value: leave-one-out MOVA value) were calculated for the test cases in each fold, and all predictions were combined into a single ROC curve. Predictions were calculated with the predict function of the randomForest package in R version 4.1.0. As with CADD, PolyPhen-2, EVE, REVEL, and AlphScore, specificity and sensitivity at each threshold were calculated with the prediction function of the ROCR package, and the ROC curve plot and AUC were calculated using the performance function. The cutoff value was set at the Youden index (the point at which sensitivity + specificity −1 is the maximum value), and sensitivity, specificity, accuracy, and precision were calculated at that cutoff value.

### Combination with other missense prediction scores

MOVA was combined with REVEL or CADD using logistic regression as implemented in the R-function glm with the option family=binomial in R version 4.1.0 and evaluated using the leave-one-out cross-validation method. The combination of AlphScore with REVEL or CADD was used directly from https://zenodo” https://zenodo.org/record/6288139#.Ym0Ir9rP23A.

### Generation of the final MOVA model

The predicted value of the probability of each variant being pathogenic was calculated by MOVA, and this was used as the MOVA value. The model was fitted using all the variants in the dataset as training data. Since machine learning can change values from each trial, the model was generated in a prediction trial of 100 times and the average of these predictions was used as the MOVA value (Figure 1C). The value was set to 0 if the allele was the same as the reference. MOVA values range from 0 to 1, with 1 having the highest probability of being pathogenic. These analyses were performed using the statistical software R version 4.1.0 (Supplementary tables 1 – 4).

### Spearman’s rank correlation coefficient

Spearman’s rank correlation coefficient was performed to observe the relationship between the AUC of MOVA and the number of variants in the dataset. The analysis was performed using the statistical software R version 4.1.0.

## Results

Figure 1 shows how MOVA was set up. Three features were used: the position of the mutant amino acid residue in the 3D structure of the wild-type protein predicted by AlphaFold2 and its pLDDT score, plus the evolutionary susceptibility to substitution between the reference and mutant amino acids obtained from the BLOSUM62 score. These data were used for training for each gene employing the random forest method. After training, the leave-one-out MOVA value (looMV) was obtained for each rare variant using the leave-one-out cross-validation method, and the AUC was calculated. For other *in silico* prediction methods, AUCs were calculated from the pre-computed predictions. In this study, an AUC of 0.70 or higher was considered a useful result.[15] The sensitivity, specificity, accuracy, and AUC for each gene in MOVA are summarized in Table 1.

**Table 1.** Performance of MOVA for the 6 ALS-causative genes.

|  | cutoff | accuracy | precision | sensitivity | specificity | AUC |
| --- | --- | --- | --- | --- | --- | --- |
| *TARDBP* | 0.612 | 0.739 | 0.8 | 0.755 | 0.714 | 0.755 |
| *FUS* | 0.369 | 0.898 | 0.844 | 0.731 | 0.954 | 0.844 |
| *SETX* | 0.042 | 0.714 | 0.083 | 0.615 | 0.718 | 0.697 |
| *TBK1* | 0.282 | 0.699 | 0.22 | 0.273 | 0.791 | 0.48 |
| *OPTN* | 0.201 | 0.784 | 0.2 | 0.333 | 0.838 | 0.438 |
| *SOD1* | 0.945 | 0.731 | 0.977 | 0.726 | 0.731 | 0.787 |

AUC : Area under the curve

MOVA showed an AUC of ≥0.70 for 3 (*TARDBP*, *FUS*, *SOD1*) of the 6 genes (Figure 2A). In PolyPhen-2, only one *(TBK1)* of the six genes showed an AUC of ≥0.70. In EVE, 2 (*OPTN*, *SOD1*) of the 4 genes (*TARDBP*, *SETX*, *OPTN*, *SOD1*) listed in the database (https://evemodel“ https://evemodel.org/) showed an AUC of ≥0.70. In AlphScore and CADD, 2 (*TBK1*, *SOD1*) of the 6 genes, and in REVEL 5 genes (*TARDBP*, *FUS*, *TBK1*, *OPTN*, *SOD1*) showed an AUC of ≥0.70. *SETX* did not show an AUC of ≥0.70 for any of the methods.

**Figure 2.**
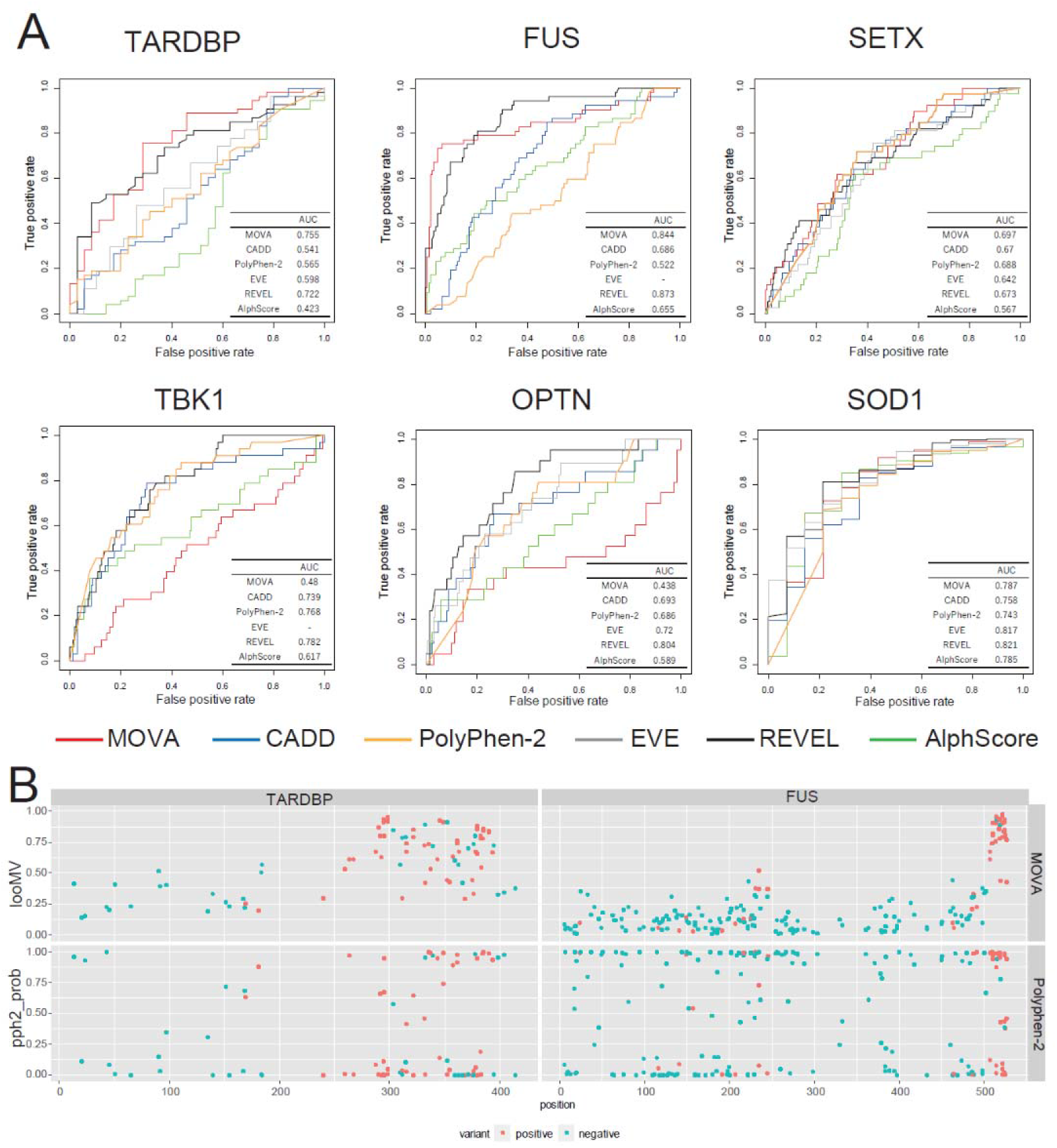
We used receiver operating characteristic (ROC) curve analysis to determine whether MOVA (red line), CADD (blue line), PolyPhen-2 (orange line), EVE (gray line), REVEL (black line), or AlphScore (green line) classified variants for *TARDBP*, *FUS*, *SETX*, *TBK1*, *OPTN*, and *SOD1* as positive or negative. The area under the ROC curve (AUC) is represented in the tables at the lower right of each graph (A). All variants of *TARDBP* and *FUS* in the dataset were divided into positive variants (red) associated with ALS and negative variants (light blue) recognized in the general population. For Polyphen-2, pph2_prob (classifier probability of the variation being damaging), and MOVA, the looMV value was plotted on the y-axis as the predicted value and the residue number was plotted on the x-axis. Both pph2_prob and looMV take values between 0 and 1, with 1 having the highest probability of being pathogenic (B).

For each gene variant, the predicted probability of pathogenicity was compared between MOVA and PolyPhen-2. For *TARDBP* and *FUS*, MOVA tended to show higher looMV values at these hotspots. In contrast, no such trend was observed for pph2_prob in Polyphen-2 (Figure 2B). MOVA values for each mutation in *TARDBP*, *FUS*, *SETX*, and *SOD1* are shown in Supplementary tables 1, 2, 3, and 4.

*SETX* was difficult to predict by any of the methods; *SETX* has ataxia/eye paralysis2 and ALS phenotypes in the same gene, which may have affected the prediction accuracy. MOVA may improve prediction accuracy for phenotypic diversity due to different mutations of the same gene by learning on a specific phenotype. Therefore, we investigated this possibility using *SETX*.[16] When both ataxia/ophthalmoplegia2 and ALS, and both associated mutations, were used as supervised data, MOVA had a lower AUC than PolyPhen-2, CADD, and REVEL (Supplementary Fig. 1). However, when only the ALS-associated mutation was used as the teacher data, MOVA yielded a higher AUC than the other methods (Figure 2A).

Finally, logistic regression analysis was used to examine whether combining MOVA with CADD or REVEL could improve prediction accuracy. The results showed that combining MOVA and CADD improved prediction accuracy, resulting in an improvement relative to CADD alone for *TARDBP* and *FUS*, with an of AUC ≥0.70 for four genes (Supplementary Fig. 2). These results were better than those for CADD combined with AlphScore; REVEL combined with MOVA yielded an AUC similar to that for REVEL alone (Supplementary Figure 3).

## Discussion

A new *in silico* method for predicting the pathogenicity of missense variants, MOVA, was developed by machine learning using the pathogenicity information previously reported for each gene as training data, using the position of the mutation site on the 3D structure of each protein predicted by Alphafold2 as the feature value. Similarly, we compared the prediction accuracy of MOVA with that of AlphScore, which predicts pathogenicity using the three-dimensional structural data of AlphaFold2. When used for analysis of six ALS-causative genes, MOVA showed higher prediction accuracy than AlphScore for four of them (*TARDBP*, *FUS*, *SETX*, and *SOD1*). Especially for *TARDBP* and *FUS*, MOVA was superior for determining pathogenicity at hotspots; MOVA performs machine learning on a gene-by-gene basis, which may explain its superior performance for predicting the pathogenicity of polymorphisms in pathogenic genes with hotspots.

Since this method focuses on specific genes for learning, we compared the prediction accuracy of MOVA with five existing methods (CADD, polyphen2, EVE, REVEL, and Alphscore) for the six ALS-causative genes. Of these, REVEL performed best for all genes. MOVA performed almost as well as REVEL, except for *TBK1* and *OPTN*. In particular, only MOVA and REVEL achieved an AUC of ≥0.7 for *TARDBP* and *FUS*. AlphScore increased prediction accuracy when combined with CADD or REVEL. MOVA, when combined with CADD or REVEL, showed a similar or better discrimination rate than AlphScore. These results demonstrate the utility of MOVA for predicting the pathogenicity of ALS-causative variant genes.

With regard to the characteristics of genes for which MOVA is effective, it is a machine-learning program that considers a variant’s position in the 3D structure predicted by AlphaFold2, the pLDDT score, which is the certainty of the prediction,[11] and the likelihood of evolutionary change between the reference and the alternative amino acid residue derived from the BLOSUM62 score, using previously reported pathogenic mutation information as the teacher data. Therefore, it is effective when the pathogenicity depends on the position of the variant in the 3D structure and its localization to a specific region. If there is a hot spot of genetic variation, MOVA may increase the prediction accuracy. Indeed, in *TARDBP* and *FUS*, for which MOVA was effective, pathogenic mutations are concentrated at the C-terminus, and moreover, in *TARDBP*, they are concentrated in IDRs (intrinsic disordered regions) (Figure 2 B, red circles).[9, 10] Since pLDDT scores are associated with IDRs,[17] MOVA may also be superior for detecting genes that are associated with IDRs and pathogenicity. However, in *TBK1* or *OPTN*, where the prediction accuracy of MOVA was low, there are no regions of concentrated pathogenic mutations,[18, 19] suggesting that the characteristics of MOVA may not have been exploited.

The prediction accuracy of machine learning is known to depend on the number of teacher data. In this study, two types of teacher data were used: pathogenic and neutral mutants. More than 50 pathogenic mutants were used for *TARDBP*, *FUS*, and *SOD1*, for which MOVA showed high prediction accuracy. On the other hand, 33 and 21 were used for *TBK1* and *OPTN*, respectively, which had low prediction accuracy. Spearman’s coefficient of rank correlation between the number of pathogenic variants used in the analysis and the MOVA AUC for each gene was 0.83. Although the number of genes analyzed was small and no significant correlation was obtained (p = 0.058), the number of pathogenic variants used as teacher data in MOVA may have affected the prediction accuracy. However, the number of neutral mutations was more than 200 for *FUS*, *SETX*, *OPTN*, and *TBK1*, and less than 100 for *TARDBP* and *SOD1*; no association with MOVA prediction accuracy was found. Statistical analysis showed that the coefficient of Spearman’s rank correlation between the number of neutral variants used in the analysis and MOVA AUC was −0.55 (p = 0.26).

MOVA performs machine learning using teacher data for each gene. Therefore, even if a gene causes different phenotypes through its mutations, it is possible to predict phenotype-specific pathogenicity by considering only those mutations that cause a particular phenotype as pathogenic mutations. In fact, using only the ALS-related gene mutations in *SETX* as the teacher data, we obtained somewhat better results than with other analytical methods in terms of the accuracy for predicting the pathogenicity of rare variants related to ALS.

One limitation of MOVA is that its prediction accuracy declines when there are limited teacher data. Additionally, it cannot be used for genes for which no teacher data are available, such as newly identified disease-related genes. Finally, only ALS-related genes were considered in this study. It remains to be seen whether other genes will show the same trend as those investigated in this study.

We have thus developed MOVA, a machine learning approach using the 3D structural position of a variant, as predicted by AlphaFold2, for learning and predicting whether or not a specific gene variant will be pathogenic. This method showed a discrimination rate for pathogenicity comparable to that of REVEL, which has the highest prediction accuracy, except for the *TBK1* and *OPTN* genes. It also showed a high pathogenicity discrimination rate when combined with CADD and REVEL. This method will be useful for predicting the pathogenicity of rare gene variants in which mutations are clustered at specific structural sites.

## Contributors

Study concept and design: YH, TI and OO. Database searches and data extraction: YH. Analysis: YH. Drafting of the manuscript: YH, TI and OO. All authors approved the final version.

## Study Funding

This research was supported through a Grant-in-Aid from the Tsubaki Memorial Foundation and SERIKA Foundation (to Y.H.) and a grant-in-aid for Scientific Research (21K07272) from the Japan Society for the Promotion of Science (to T.I).

## Competing Interests

There are no associations with companies or organizations that would constitute a conflict of interest requiring disclosure in relation to this study.

## Patient consent for publication

Not applicable.

## Ethical approval

This study did not involve humans or animals.

## Source code

The source code used in this study is available at https://github.com/yuya-hatano/MOVA.

## Figure legends

Supplementary Figure 1 We used receiver operating characteristic (ROC) curve analysis to determine whether MOVA (red line), CADD (blue line), Polyphen-2 (orange line), EVE (gray line), REVEL (black line), or AlphScore (green line) classified variants for *SETX* as positive and negative. Unlike Figure 2, all phenotypic variants were included as positive variants in both the training and validation data. The area under the ROC curve (AUC) is shown in the table at the bottom right of each graph (A).

Supplementary Figure 2 We used receiver operating characteristic (ROC) curve analysis to determine whether MOVA + CADD (red line), MOVA (blue line), CADD (black line), or CADD + AlphScore (gray line) classified variants for *TARDBP*, *FUS*, *SETX*, *TBK1*, *OPTN*, and *SOD1* as positive and negative. The area under the ROC curve (AUC) is represented in the tables at the lower right of each graph.

Supplementary Figure 3 We used receiver operating characteristic (ROC) curve analysis to determine whether MOVA + REVEL (red line), MOVA (blue line), REVEL (black line), or REVEL + AlphScore (gray line) classified variants for *TARDBP*, *FUS*, *SETX*, *TBK1*, *OPTN*, and *SOD1* as positive and negative. The area under the ROC curve (AUC) is represented in the tables at the lower right of each graph.

Supplementary Table 1

MOVA values for each variant in *TARDBP*.

Ref: reference allele, Pos,: position alt: alternative allele.

Supplementary Table 2

MOVA values for each variant in *FUS*.

Ref: reference allele, Pos,: position alt: alternative allele.

Supplementary Table 3

MOVA values for each variant in *SETX*.

Ref: reference allele Pos,: position alt: alternative allele.

Supplementary Table 4

MOVA values for each variant in *SOD1*.

Ref: reference allele, Pos,: position alt: alternative allele.

## Notes

### Competing Interest Statement

The authors have declared no competing interest.

