## Supplementary figures and images for "Accuracy of a machine learning method based on structural and locational information from AlphaFold2 for predicting the pathogenicity of *TARDBP* and *FUS* gene variants in ALS"

### Supplementary Figure 1

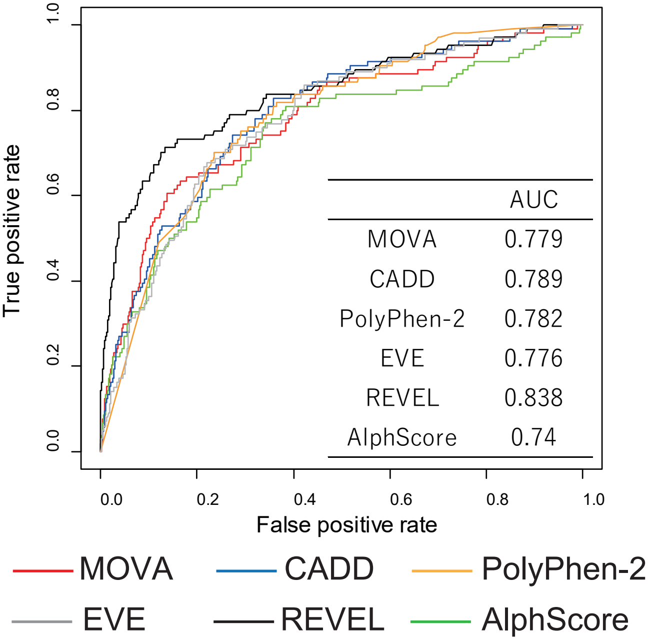

### Supplementary Figure 2

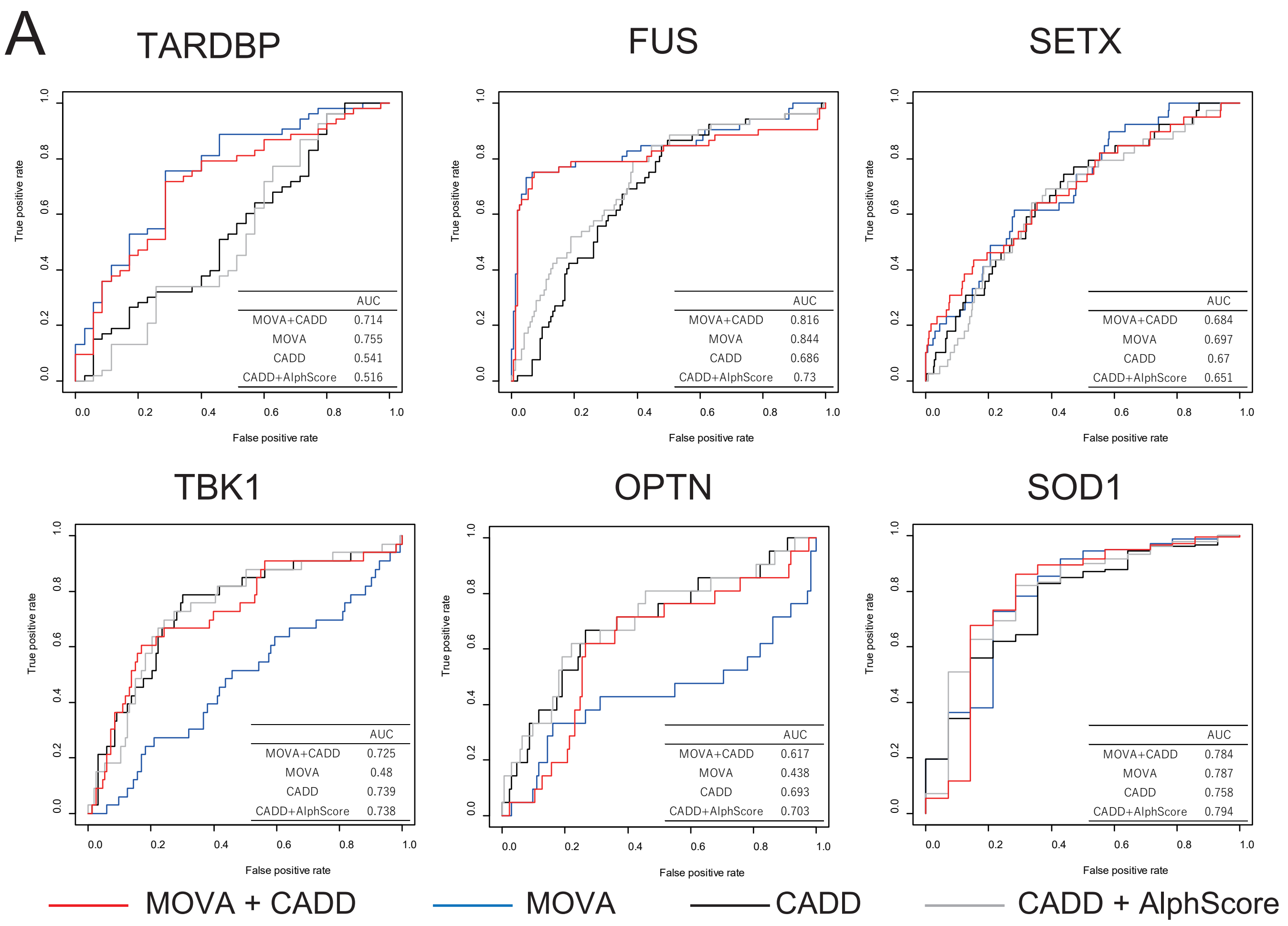

### Supplementary Figure 3

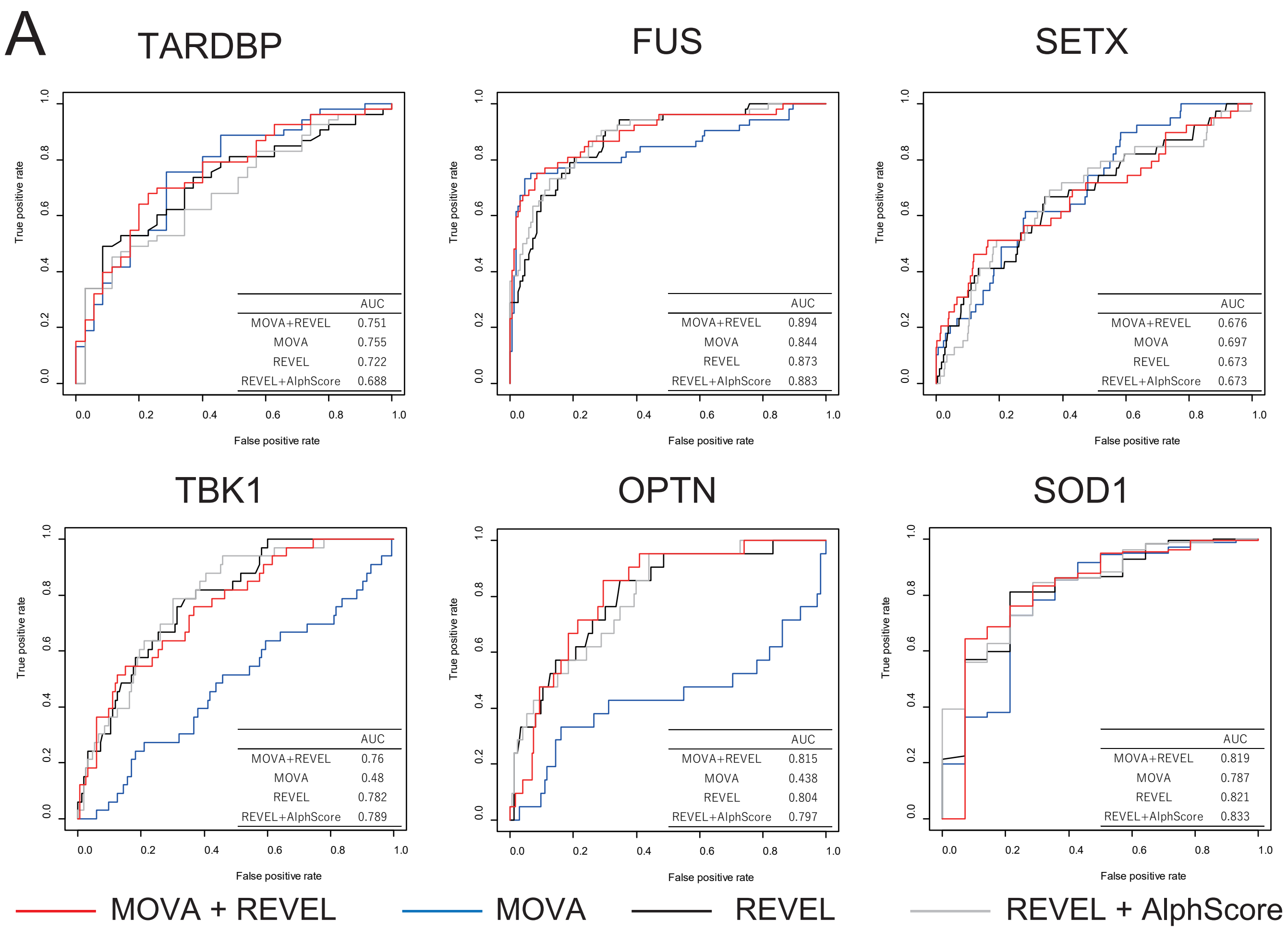
